# *Plasmodium simium* causing human malaria: a zoonosis with outbreak potential in the Rio de Janeiro Brazilian Atlantic forest

**DOI:** 10.1101/122127

**Authors:** Patrícia Brasil, Mariano Gustavo Zalis, Anielle de Pina-Costa, Andre Machado Siqueira, Cesare Bianco Júnior, Sidnei Silva, André Luiz Lisboa Areas, Marcelo Pelajo-Machado, Denise Anete Madureira de Alvarenga, Ana Carolina Faria da Silva Santelli, Hermano Gomes Albuquerque, Pedro Cravo, Filipe Vieira Santos de Abreu, Cassio Leonel Peterka, Graziela Maria Zanini, Martha Cecilia Suárez Mutis, Alcides Pissinatti, Ricardo Lourenço-de-Oliveira, Cristiana Ferreira Alves de Brito, Maria de Fátima Ferreira-da-Cruz, Richard Culleton, Cláudio Tadeu Daniel-Ribeiro

**Affiliations:** Laboratório de Doenças Febris Agudas, Instituto Nacional de Infectologia Evandro Chagas (INI), Fundação Oswaldo Cruz (Fiocruz), Rio de Janeiro, Brazil; Centro de Pesquisa, Diagnóstico e Treinamento em Malária (CPD-Mal), Fiocruz, Rio de Janeiro, Brazil; Laboratório de Infectologia e Parasitologia Molecular, Hospital Universitário Clementino Fraga Filho, Universidade Federal do Rio de Janeiro, Rio de Janeiro, Brazil; Laboratório de Pesquisa em Malária, Instituto Oswaldo Cruz (IOC), Fiocruz, Rio de Janeiro, Brazil; Centro Universitário Serra dos Órgãos (UNIFESO), Teresópolis, Rio de Janeiro, Brazil; Laboratório de Parasitologia, INI, Fiocruz, Rio de Janeiro, Brazil; Laboratório de Patologia, IOC, Fiocruz, Rio de Janeiro, Brazil; Laboratório de Malária, Centro de Pesquisas René Rachou (CPqRR), Fiocruz, Belo Horizonte, Brazil; Programa Nacional de Prevenção e Controle da Malária, Secretaria de Vigilância em Saúde, Ministério da Saúde, Brasília, DF, Brazil; Laboratório de Doenças Parasitárias, IOC, Fiocruz, Rio de Janeiro, Brazil; Laboratório de Genoma e Biotecnologia (GenoBio), Instituto de Patologia Tropical e Saúde Pública, Universidade Federal de Goiás, Goiânia, Brazil; Instituto de Higiene e Medicina Tropical, Universidade Nova de Lisboa, Lisbon, Portugal; Laboratório de Mosquitos Transmissores de Hematozoários, IOC, Fiocruz, Rio de Janeiro, Brazil; Centro de Primatologia do Rio de Janeiro (CPRJ/INEA), Brazil; Malaria Unit, Department of Pathology, Institute of Tropical Medicine, Nagasaki University, Nagasaki, Japan

## Abstract

**Background:** Malaria was eliminated from Southern and Southeastern Brazil over 50 years ago. However, an increasing number of autochthonous episodes attributed to *Plasmodium vivax* have been recently reported in the Atlantic forest region of Rio de Janeiro State. *As P. vivax*-like non-human primate malaria parasite species *Plasmodium simium* is locally enzootic, we performed a molecular epidemiological investigation in order to determine whether zoonotic malaria transmission is occurring.

**Methods:** Blood samples of humans presenting signs and/or symptoms suggestive of malaria as well as from local howler-monkeys were examined by microscopy and PCR. Additionally, a molecular assay based on sequencing of the parasite mitochondrial genome was developed to distinguish between *P. vivax* and *P. simium*, and applied to 33 cases from outbreaks occurred in 2015 and 2016.

**Results:** Of 28 samples for which the assay was successfully performed, all were shown to be *P. simium*, indicating the zoonotic transmission of this species to humans in this region. Sequencing of the whole mitochondrial genome of three of these cases showed that *P. simium* is most closely related to *P. vivax* parasites from South American.

**Findings:** The explored malaria outbreaks were caused by *P. simium*, previously considered a monkey-specific malaria parasite, related to but distinct from *P. vivax*, and which has never conclusively been shown to infect humans before.

**Interpretation:** This unequivocal demonstration of zoonotic transmission, 50 years after the only previous report of *P. simium* in man, leads to the possibility that this parasite has always infected humans in this region, but that it has been consistently misdiagnosed as *P. vivax* due to a lack of molecular typing techniques. Thorough screening of the local non-human primate and anophelines is required to evaluate the extent of this newly recognized zoonotic threat to public health and malaria eradication in Brazil.

**Funding:** Fundação Carlos Chagas Filho de Amparo à Pesquisa do Estado de Rio de Janeiro (Faperj), The Brazilian National Council for Scientific and Technological Development (CNPq), JSPS Grant-in-Aid for scientific research, Secretary for Health Surveillance (SVS) of the Ministry of Health, Global Fund, and PRONEX Program of the CNPq.

## Introduction

Zoonotic malaria occurs when humans become infected with malaria parasite species that more commonly infect non-human primates. Species such as *Plasmodium knowlesi* and *Plasmodium cynomolgi*, both parasites of macaque monkeys, can infect humans *via* the bites of infected mosquitoes under natural and experimental conditions. Although no transmission from humans to mosquitoes has yet been documented in the field, *P. knowlesi* has recently been shown to be responsible for many cases of human malaria in Southeast Asia, mostly affecting individuals living or working in close contact with forests.^1^ Zoonotic malaria poses a unique problem for malaria control efforts and complicates the drive towards eventual elimination of the disease.

Once prevalent throughout the country, malaria transmission in Brazil currently occurs almost entirely within the northern Amazon region of the country. However, there are a consistent number of autochthonous cases reported in southern and southeastern Brazil; regions from where human malaria was eliminated 50 years ago.^2^ Most of these episodes are attributed to *P. vivax*, predominantly occur in areas of dense Atlantic forest vegetation, and mainly affect non-resident visitors. In most outbreaks, there is no identifiable index case that could have introduced the parasite from a malaria endemic region.^3^

It has been hypothesized that autochthonous “*P. vivax”* malaria in the Atlantic forest could be the result of infection by malaria parasites species that commonly infect non-human primates^4^. Deane^5^ proposed that monkeys could serve as reservoirs of *Plasmodium* that could infect humans. In the forests of continental America, two species of *Plasmodium* infecting non-human primates have been described: *Plasmodium simium* Fonseca, 1951 and *Plasmodium brasilianum* Gonder & von Berenberg-Gossler, 1908. These are similar at the morphological, genetic, and immunological levels to *P. vivax* and *P. malariae*, respectively.^5^-^9^

In the state of Rio de Janeiro, located along the coast in the Southeast region of Brazil, sporadic autochthonous malaria cases have been reported from the Atlantic forest area since 1993. Between 2006 and 2014, sixty-one autochthonous malaria cases were reported in Rio de Janeiro, an average of 6·8 cases per year, with an unexpectedly high incidence in 2015 (n=33) and 2016 (n=16).

Here we describe the parasitological and molecular analyses of parasites causing autochthonous human malaria in the Atlantic forest region of Rio de Janeiro in 2015 and 2016 with the aim of determining whether zoonotic malaria transmission occurs there.

## Methods

### Study area and population

Rio de Janeiro State is located in the Southeast region of Brazil. It is essentially a mosaic of urban areas with high population densities, mostly in the coastal lowlands, and mountain areas covered by the Atlantic forest containing small cities and settlements scattered in the valleys.^10^ Localities where malaria cases have been reported are situated in valleys between 280 and 1300 meters above sea level.^11^ Detailed descriptions of the environmental characteristics of the Atlantic forest are published elsewhere.^3^

Cases considered here are from patients who attended the *Instituto Nacional de Infectologia Evandro Chagas* (INI-IPEC), a reference centre for the diagnosis and treatment of infectious diseases at Fiocruz, in Rio de Janeiro. Blood samples from patients suffering from acute fever symptoms were collected from the Acute Febrile Illness Outpatient Clinic (*Ambulatório de Doenças Febris Agudas, DFA*).

### Ethical clearance

The INI-Fiocruz Ethical Committee in Research approved the study (N. 0062.0.009.000-11). All participants provided informed consent.

### Study design

This is a descriptive study with parasitological and molecular characterization of the autochthonous malaria cases acquired in the Rio de Janeiro Atlantic forest in 2015 and 2016. An epidemiological investigation was performed to characterize the likely location of infection for classifying each episode as autochthonous.

### Patient Procedures

Individuals were recruited upon presentation of signs and/or symptoms suggestive of malaria, a history of travel to or habitation in areas within the Rio de Janeiro Atlantic forest and a positive test by blood smear and/or PCR. The following exclusion criteria were applied: malaria prophylaxis, blood transfusion or tissue/organ transplantation, use of intravenous drugs, needle stick injury, residence or recreation near ports or airports or travel to known malaria endemic areas in the preceding year. Following informed consent, venous blood was drawn for clinical laboratory analyses and molecular studies. Additional tests such as blood culture and viral serology were performed at the attending physician’s discretion.

### Diagnosis by microscopy

Giemsa-stained thin and thick blood smears were examined by bright-field microscopy, using a 100x/1·3 NA oil immersion objective lens for species identification and parasite density estimation.^12^ Blood films were examined for a minimum of 100 fields in the case of the presence of malaria parasites and 500 fields when no parasites could be detected. Parasite numbers were recorded per 200 white blood cells (WBC) or 500 WBCs in the case of low parasitemia. To estimate parasite density per μL of blood, a standard mean WBC count of 6,000/ μL blood was assumed. All slides were subsequently examined by an independent malaria microscopist qualified by PAHO/WHO malaria accreditation course.

### DNA extraction and PCR

DNA was extracted from whole blood with the QIAamp midi kit^®^, according to the manufacturer’s protocol (Qiagen). DNA samples were tested for *P. vivax* by conventional PCR^13^ and by real time PCR, both using the cysteine-proteinase gene as a target (GenBank number L26362). For real time PCR, 2·5 μl of DNA were added to a 47·5 μL mixture containing the 1× TaqMan^®^ Universal PCR Master Mix (Applied Biosystems), 300 nM of each primer Pv1 (5’ATCAACGAGCAGATGGAGAAATATA3’) and Pv5 (5’GCTCTCGAAATCTTTCTTCGA3’), and 150 nM of PVIV probe (5’ FAM^™^ AACTTCAAAATGAATTATCTC MGB^™^ NFQ 3’) (Applied Biosystems). Amplification conditions involved two holds (50 °C/2 minutes and 95 °C/10 minutes) followed by 40 cycles of amplification (95 °C/15 seconds and 60 °C/1 minutes). The real-time PCR was run at least in duplicate on the ABI PRISM Sequence Detection System 7500 (Applied Biosystems). The results were analyzed using the real-time software (ABI Prism 7500 software v. 1.1 RQ study). TaqMan^®^ RNaseP Control 20x (Applied Biosystems, California, EUA) was used as endogenous reaction control. To avoid DNA contamination, separated workstations for mix preparation and DNA extraction were utilized and DNA Away^®^ was applied for surface decontamination. Positive (DNA extracted from blood from patients with known *P. vivax* infection) and negative (no DNA and DNA extracted from individuals who have never traveled to malaria-endemic areas) controls were used in each round of amplification.

### Non-human primate samples

DNA was extracted from samples obtained from four howler monkeys from distinct sites and times in Southeastern Brazil. DNA extracted from the spleen and liver of one brown howler monkey (*Alouatta clamitans*) found dead in Guapimirim (one of the municipalities where human malaria occurs in the Rio de Janeiro Atlantic forest), was used for *Plasmodium* detection by nested-PCR.^14^ Samples from both organs were positive for *P. vivax/P.simium* DNA.^15^ DNA was also extracted from blood of two *A. clamitans*, one of them captured in 2016 at Vale das Princesas, Miguel Pereira (a site where human cases have also been reported in the Rio de Janeiro state), which tested positive for both *P. vivax/P. simium* and *P. brasilianum/malariae*, and one other from Macaé (another locality with human cases), also positive for *P. vivax/P. simium*. In addition, a *P. simium* reference sample (ATCC^®^ 30130^™^), derived from a howler monkey (*Alouatta fusca*) captured in São Paulo, Southeast Brazil, in 1966, was also used(https://www.atcc.org/∼/ps/30130.ashx).

DNA was extracted from a sample acquired from ATCC in 1994 and kept frozen in N2 until use. The DNA extracted from all these four monkey samples was further submitted to mitochondrial genome analysis.

### Molecular phylogenetic analysis of *P. simium* infections

Thirty-three samples were obtained from autochthonous human cases, two from monkeys of Rio de Janeiro Atlantic forest transmission sites and one ATCC *P. simium* reference sample and were subjected to malaria parasite mitochondrial genome sequencing.

Whole mitochondrial genome sequencing was performed on a subset of the human infections (three cases: AF1, AF2, and AF3) and the ATCC reference sample, following the methodology described in Culleton *et al* (2011) and compared with 794 *P. vivax* mitochondrial genome sequences and three sequences of *P. simium* (accession numbers AY800110, NC_007233 and AY722798, all of which have identical sequences) deposited in Genbank.^16-22^ Using these sequences, a median-joining haplotype network was produced with NETWORK 4.5.0 (Fluxus technology Ltd. 2006), as previously described.^23^

The mitochondrial genome of the other thirty samples was partially sequenced to distinguish *P. simium* samples from *P. vivax. Plasmodium simium* differs from the most closely related *P. vivax* isolate at two unique single nucleotide polymorphisms (SNPs) in the mitochondrial genome, at positions 3535 (T−>C) and 3869 (A−>G), numbered according to the nucleotide sequences deposited by Culleton *et al* (2011).^23^ These two SNPs are close together, and can be PCR amplified and sequenced with a single set of primers, or with a nested PCR if DNA concentrations are low. Primer pairs for the outer PCR are Outer PCR Primers: *PsimOUTF* 5’CAGGTGGTGTTTTAATGTTATTATCAG3’ and *PsimOUTR* 5’CATAGGTAAGAATGTTAATACAACTCC3’. For the inner PCR, the primers are *PsimINF* 5’GCTGGAGATCCTATTTTATATCAAC3’ and *PsimINR* 5’ATGTAAACAATCCAATAATTGCACC3’.

## RESULTS

In 2015, 33 autochthonous malaria cases occurred in the Rio de Janeiro state, 25 (76%) of them processed at the DFA outpatient clinic of Fiocruz. In 2016 (up to October 31^st^), 16 autochthonous malaria cases occurred in Rio de Janeiro, 14 (87·5%) seen at the DFA-Fiocruz, totaling 39 (79·6%) out of the 49 cases reported in the state. In all cases, transmission occurred in low-population density non-urban sites located in Atlantic forest areas (Figure 1). It was the first malaria episode for all patients, and none presented clinical or laboratory complications associated with malaria. In 37 cases the diagnosis of *P. vivax* was made by microscopy. The highest parasitaemia was 3,000 parasites per μL of blood and, in more than 63% of the cases, it was lower than 500 parasites per μL. Two patients were negative for the presence of parasites by microscopy. A PCR assay for *P. vivax* species detection, but which does not discriminate between *P. vivax* and *P. simium*, suggested the presence of *P. vivax* in 38 (out of the 39) patients.

**Figure 1:**
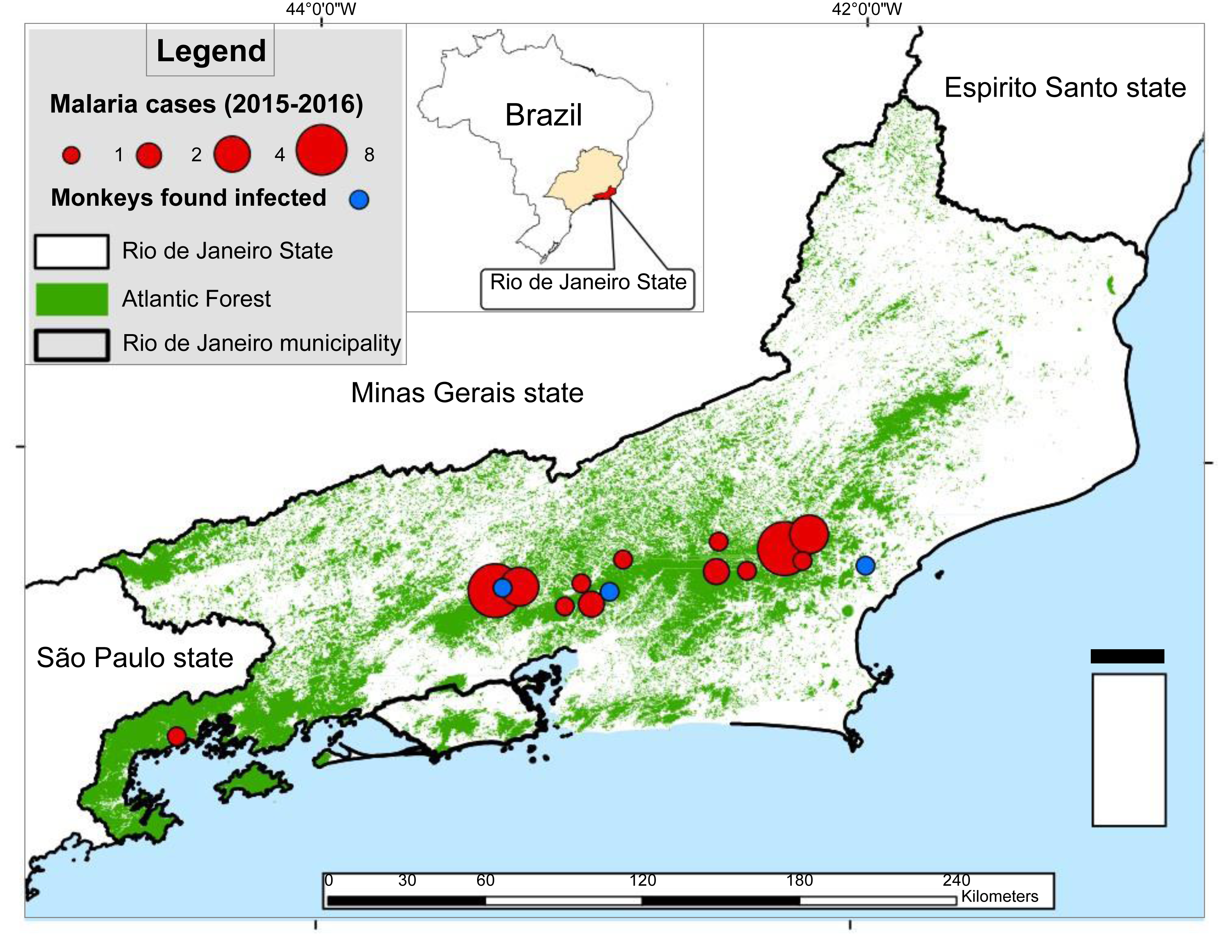
Map of the Rio de Janeiro state showing the Atlantic Forest and indicating where human malaria cases of simian origin and *P. simium* infected monkeys have been detected. Human cases are represented by red spots of different sizes (symbolizing one to eight cases), and the three captured infected wild howler monkeys are shown as blue spots. The extension of the area covered by the Atlantic forest vegetation is indicated in green. All cases were reported in forest fragments located in the Serra do Mar and a monkey carrying *P. simium* was found in the vicinity of each area. The municipality of Rio de Janeiro, delimitated in bold line, is free of malaria transmission.

### Morphological characteristics

When compared with *P. vivax* from the malaria endemic Amazonian regions, parasites from the Atlantic forest diagnosed as *P. vivax* were morphologically different (Suppl. Table 1). Trophozoites are pleomorphic but less amoeboid than those observed in *P. vivax*. They had a large mass of chromatin and a more compact cytoplasm with malaria pigment. Usually stippling was mostly observed in infected cells with late developmental forms, but erythrocytes containing early trophoizoites were also frequently dotted (Figure 2, A-F). In addition, developing schizonts contained fewer merozoites than in *P. vivax* (Figure 2, G-L). The highest number of merozoites in mature schizonts was 14 (Figure 2, M). Gametocytes were round with compact cytoplasm and marked pigmentation (Figure 2, N-P).

**Figure 2:**
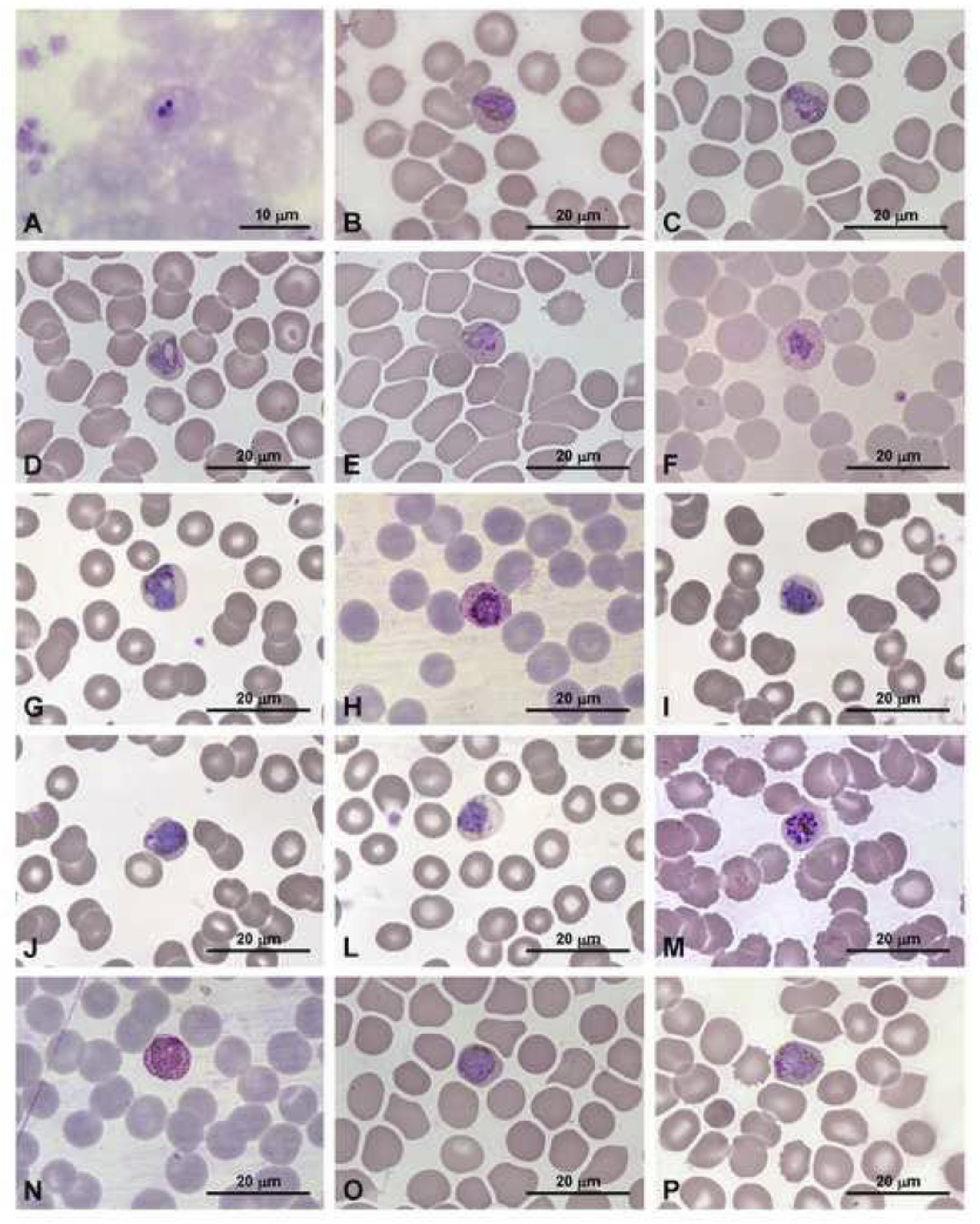
Giemsa-stained preparations of blood samples from humans naturally infected with *Plasmodium simium* in Rio de Janeiro state, Brazil. All preparations are thin blood-films, except A (thick blood smear). A: early throphozoite; B-F: pleomorphic developing trophozoites; G-L: immature schizonts; M: mature schizont; N-P: gametocytes.

Non-infected erythrocytes showed marked anisocytosis and poikilocytosis. The latter was represented mainly by acanthocytes, dacrocytes and spherocytes, which occurred together on the same preparations.

### Molecular Characterization

Twenty samples from 25 cases occurring in 2015 and 13 samples from the 14 cases in 2016 were analyzed by molecular tests in view of the diagnosis of the *Plasmodium* sp. Because of the low amount and bad quality of extracted DNA, the full-length mitochondrial genome sequence was obtained for only four samples (three human samples and one *P. simium* reference strain).

Analysis of this mitochondrial genome sequence revealed that all four samples shared identical sequences, and these were identical to the mitochondrial genome sequence of *P. simium* available at genbank, which differs from the most closely related isolates of *P. vivax* by two unique *P. simium*-specific SNPs. Analysis of 794 full-length mitochondrial genome sequences from globally acquired *P. vivax* samples^1-7^ confirmed that these SNPs were unique to *P. simium*. A haplotype network tree (Figure 3) was constructed using these sequences, and clearly shows that *P. simium* is most closely related to the *P. vivax* parasites of man isolated from South America.

**Figure 3:**
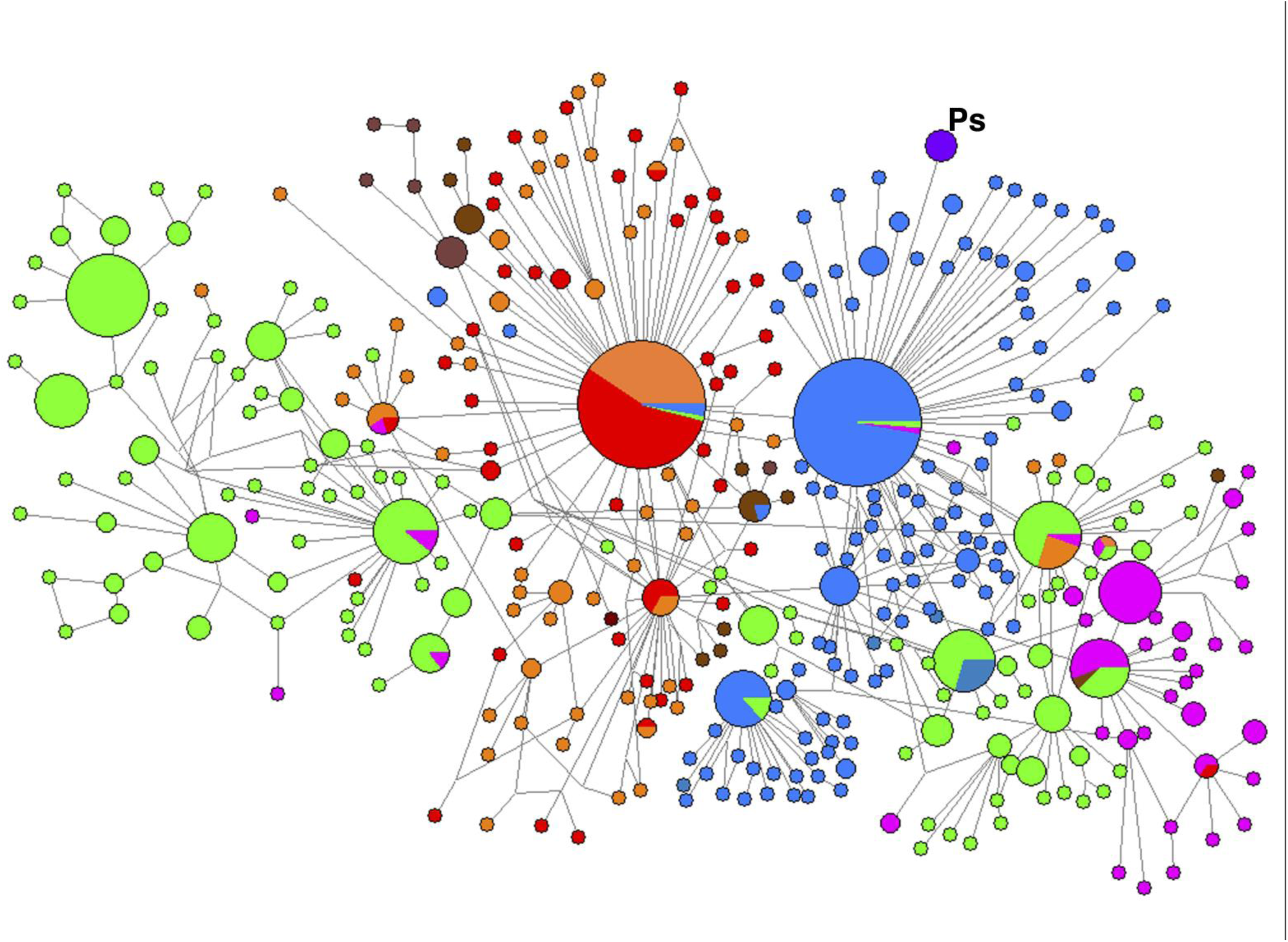
Haplotype network constructed using NETWORK 4.5.0. It is incorporating 794 *Plasmodium vivax* mitochondrial genome sequences (from References^1-7^), three *Plasmodium simium* sequences (Accession numbers AY800110, NC_007233 and AY722798), and three sequences from the current outbreak in Rio de Janeiro state (sample numbers AF1, AF2 and AF3). The *P. simium* samples and those from the current outbreak share an identical haplotype, colored purple and marked “Ps”, and are distinct from all other isolates. Node sizes are proportional to haplotype frequency. Node colours indicate the geographic origin of the isolates and are coded as follows; red, Africa; blue, South America; green, Asia; pink, Melanesia; orange, the Indian sub-continent; brown, Middle East (Turkey and Iran).

### Characterization of further infections using informative mtDNA SNPs

Of 33 samples species-typed based on the two informative SNPs that differentiate *P. vivax* from *P. simium*, we were able to diagnose an infection of *P. simium* in 28 of them (Table 1). We were unable to achieve PCR amplification for the remaining five samples, due to technical constraints. The same informative SNPs were found in *Plasmodium simium* infecting three local howler monkeys, MB CPRJ, RJ30 and RJ59 (Table 1).

**Table 1:** Identification of *Plasmodium simium* in human and simian samples through partial or whole mitochondrial genome sequencing.

| *HUMAN SAMPLES* |  |  |  |
| --- | --- | --- | --- |
| Sample |  | *P. simium* SNP | *P. simium* |
| Identification | Year | Sequencing | Whole mitochondrial sequencing |
| AF 1 | 2015 | *P. simium* | *P. simium* |
| AF 2 | 2015 | *P. simium* | *P. simium* |
| AF 3 | 2015 | *P. simium* | *P. simium* |
| AF 4 | 2015 | *P. simium* | - |
| AF 5 | 2015 | *P. simium* | - |
| AF 6 | 2015 | *P. simium* | - |
| AF 7 | 2015 | *P. simium* | - |
| AF 8 | 2015 | *P. simium* | - |
| AF 9 | 2015 | *P. simium* | - |
| AF 10 | 2015 | *P. simium* | - |
| AF 11 | 2015 | *P. simium* | - |
| AF 12 | 2015 | *P. simium* | - |
| AF 13 | 2015 | *P. simium* | - |
| AF 14 | 2015 | Not amplified | - |
| AF 15 | 2015 | *P. simium* | - |
| AF 16 | 2015 | *P. simium* | - |
| AF 17 | 2015 | Not amplified | - |
| AF 18 | 2015 | *P. simium* | - |
| AF 19 | 2015 | *P. simium* | - |
| AF 20 | 2015 | Not amplified | - |
| AF 21 | 2016 | *P. simium* | - |
| AF 22 | 2016 | *P. simium* | - |
| AF 23 | 2016 | *P. simium* | - |
| AF 24 | 2016 | *P. simium* | - |
| AF 25 | 2016 | *P. simium* | - |
| AF 26 | 2016 | *P. simium* | - |
| AF 27 | 2016 | *P. simium* | - |
| AF 28 | 2016 | *P. simium* | - |
| AF 29 | 2016 | Not amplified | - |
| AF 30 | 2016 | *P. simium* | - |
| AF 31 | 2016 | *P. simium* | - |
| AF 32 | 2016 | *P. simium* | - |
| AF 33 | 2016 | Not amplified | - |
| *MONKEY SAMPLES* |  |  |  |
| ATCC®30130™ | 1966 | *P. simium* | *P. simium* |
| MB CPRJ | 2013 | *P. simium* | - |
| RJ 30 | 2016 | *P. simium* | - |
| RJ 59 | 2016 | *P. simium* | - |

## Discussion

The New World non-human primate tertian malaria parasite *P. simium* was first identified by Fonseca (1951) in a monkey from the state of São Paulo, Brazil. It appears to be restricted to the Atlantic forest in southern and southeastern Brazil.^4,5,24-26^ Here, we show that *P. simium* is infecting and causing disease in humans. Fonseca (1951)^27^, Garnham (1966)^28^ and Deane highlighted the morphological differences between *P. vivax* and *P. simium;* the trophozoites of the latter being less amoeboid and with coarser and more precocious and very prominent Schuffner’s dots than *P. vivax*. Garnham (1966)^28^ pointed that the detection of stippling in *P. simium* early parasitized cells depends on the staining procedures. These morphological characteristics of *P. simium* are consistent with those described here for the infections of humans from the Atlantic forest (Figure 2, Suppl Table).

Despite initial diagnoses of *P. vivax* for these infections, molecular evidence has revealed that these parasites are *P. simium*. This misdiagnosis of a zoonotic non-human primate malaria parasite as a human parasite species has precedent and parallels the discovery of the large focus of *P. knowlesi* in Borneo, which was initially attributed to *P. malariae* on the basis of morphological characteristics.^29^

Notwithstanding the apparent genetic similarity of *P. simium* to *P. vivax*, attempts at inducing infections of *P. simium* in humans under laboratory conditions have been unsuccessful.^6^ In 1966, however, Deane *et al* described the infection of a man with a *P. vivax*-like parasite that they considered to be *P. simium*, based on morphological characteristics of the parasite and on the fact that infection had occurred in a forest reserve outside São Paulo, where *P. simium* was known to be transmitted.^30^ This remains the only previous case report of a possible human infection with *P. simium*.

We show here that *P. simium* can and does infect humans under natural conditions and can cause outbreaks of zoonotic disease in the Atlantic forest region in Rio de Janeiro. This situation has immediate implications for public health in this region, and further and more profound consequences for the control and eventual elimination of malaria in Brazil.

Patients who were naturally infected with *P. simium* reported clinical symptoms congruent to those of vivax malaria, and responded successfully to chloroquine and primaquine, with no deaths reported. It is not currently known whether *P. simium* is capable of producing hypnozoites in humans and, thus, relapses, as does *P. vivax*. However, one patient (from February 2015, AF3), who were treated solely with chloroquine due to G6PD deficiency, and one other (January 2016, AF21) who discontinued primaquine treatment due to adverse events, have not presented any relapse to date (March 2017), suggesting that hypnozoites were not formed. Further follow-up of this cohort is required to monitor potential relapse in these patients.

In summary, we report here that the malaria outbreaks in 2015, and in 2016 in the Atlantic forest of southeastern Brazil were caused by *P. simium*, previously considered to be a monkey-specific species of malaria parasite, related to but distinct from *P. vivax*, and which has never conclusively been shown to infect humans before. Such zoonotic transmission of malaria from a monkey reservoir to man has immediate consequences for public health in this region, and, furthermore, for future attempts to control and eventually eliminate malaria in Brazil. Thorough screening of the local non-human primate and anopheline mosquito populations in this area is required to evaluate the extent of this newly recognized zoonotic threat to public health.

This unequivocal demonstration of zoonotic transmission, 50 years after the only previous report of *P. simium* in man, leads to the possibility that this parasite has always infected humans in this region, but that it has been consistently misdiagnosed as *P. vivax* due to a lack of molecular typing techniques. Alternatively, it may be the case that *P. simium* has only recently acquired the ability to frequently infect humans. This latter scenario has extremely important implications in terms of parasite/host relationships and evolution.

## ACKNOWLEDGEMENTS

To Ana Paula Barroso Teixeira de Freitas and Ana Cláudia Ribeiro Fiuza for assistance on the parasitological diagnosis, to Mrs. Aline Lavigne for conducting the PCR for *P. vivax* and to Igor José Silva for help with the photography of the slides. The authors are also grateful to Doctors Silvia Bahadian Moreira, for the facilities provided at the Primate Centre of Rio de Janeiro (CPRJ), and to Cristina Maria Giordano Dias, Head for the Management of Vector Transmitted Diseases and Zoonosis of the State of Rio de Janeiro (Coordination of Epidemiological Surveillance, Superintendence of Epidemiological and Environmental Surveillance) and her team for data on the malarious patients examined outside Fiocruz. APC is supported by a post-doctoral fellowship from the *Fundação Carlos Chagas Filho de Amparo à Pesquisa do Estado de Rio de Janeiro (Faperj)*. The Brazilian National Council for Scientific and Technological Development (*CNPq*) supports PC, RLO, CFAB, MFFC and CTDR through a Research Productivity Fellowship. RLO, MFFC and CTDR are *Cientistas do nosso Estado* from the *Faperj*. RC was supported by a JSPS Grant-in-Aid for scientific research grant number 16K21233. The study received financial support from the Secretary for Health Surveillance (SVS) of the Ministry of Health, through the Global Fund (Agreement IOC-005-Fio-13) and from the PRONEX Program of the CNPq.

## REFERENCES

1 Ahmed MA, Cox-Singh J. Plasmodium knowlesi - an emerging pathogen. Isbt Science Series 2015; 10(Suppl 1): 134–140.

2 Siqueira AM, Mesones-Lapouble O, Marchesini P, et al. *Plasmodium vivax* Landscape in Brazil: Scenario and Challenges. Am J Trop Med Hyg 2016; 95(6 Suppl): 87–96.

3 de Pina-Costa A, Brasil P, Di Santi SM, et al. Malaria in Brazil: what happens outside the Amazonian endemic region. Mem Inst Oswaldo Cruz 2014; 109: 618–33.

4 Yamasaki T, Duarte AMRC, Curado I, et al. Detection of etiological agents of malaria in howler monkeys from Atlantic Forests, rescued in regions of São Paulo city, Brazil. J Med Primatol 2011; 40: 392–400.

5 Deane LM. Simian malaria in Brazil. Mem Inst Oswaldo Cruz 1992; 87: 1–20.

6 Coatney GR. The simian malarias: zoonoses, anthroponoses, or both? Am J Trop Med Hyg 1971; 20: 795–803.

7 Cochrane AH, Barnwell JW, Collins WE, Nussenzweig RS. Monoclonal antibodies produced against sporozoites of the human parasite Plasmodium malariae abolish infectivity of sporozoites of the simian parasite Plasmodium brasilianum. Infect Immun 1985; 50: 58–61.

8 Lal AA. Circumsporozoite protein gene from Plasmodium brasilianum. Animal reservoirs for human malaria parasites? J Biol Chem 1988; 263: 5495–5498.

9 Leclerc M, Hugot J, Durand P, Renaud F. Evolutionary relationships between 15 Plasmodium species from new and old world primates (including humans): an 18S rDNA cladistic analysis. Parasitology 2004; 129: 677–84.

10 Ribeiro MC, Metzger JP, Martensen AC, Ponzoni FJ, Hirota MM. The Brazilian Atlantic Forest: How much is left, and how is the remaining forest distributed? Implications for conservation. Biological Conservation 2009; 142: 1141–1153.

11 Brasil P, Costa AP, Longo CL, Silva S, Ferreira-da-Cruz MF, Daniel-Ribeiro CT. Malaria, a difficult diagnosis in a febrile patient with sub-microscopic parasitaemia and polyclonal lymphocyte activation outside the endemic region, in Brazil. Malaria Journal 2013; 12: 402.

12 Brasil. Ministério da Saúde, Secretaria de Vigilância em Saúde. Manual de diagnóstico laboratorial da malária. Ministério da Saúde, Secretaria de Vigilância em Saúde. Brasília DF: Ministério da Saúde, 2005 112p. (Série A. Normas e Manuais Técnicos). ISBN 85-334-0974-5.

13 Torres KL, Figueiredo DV, Zalis MG, Daniel-Ribeiro CT, Alecrim W, Ferreira-da-Cruz MF. Standardization of a very specific and sensitive single PCR for detection of Plasmodium vivax in low parasitized individuals and its usefulness for screening blood donors. Parasitol Res 2006; 98(6): 519–4.

14 Snounou G, Viriyakosol S, Jarra W, Thaithong S, Brown KN. Identification of the four human malaria parasite species in field samples by the polymerase chain reaction and detection of a high prevalence of mixed infections. Mol Biochem Parasitol 1993; 58: 283–92.

15 Alvarenga DA, de Pina-Costa A, de Sousa TN, et al. Simian malaria in the Brazilian Atlantic forest: first description of natural infection of capuchin monkeys (Cebinae subfamily) by Plasmodium simium. Malaria Journal 2015; 14: 81.

16 Jongwutiwes S, Putaporntip C, Iwasaki T, Ferreira MU, Kanbara H, Hughes AL. Mitochondrial genome sequences support ancient population expansion in Plasmodium vivax. Mol Biol Evol 2005; 22(8): 1733–9.

17 Mu J, Joy DA, Duan J, et al. Host switch leads to emergence of Plasmodium vivax malaria in humans. Mol Biol Evol 2005; 22(8): 1686–93.

18 Iwagami M, Hwang SY, Fukumoto M, et al. Geographical origin of Plasmodium vivax in the Republic of Korea: haplotype network analysis based on the parasite’s mitochondrial genome. Malaria Journal 2010; 9: 184.

19 Culleton R, Coban C, Zeyrek FY, et al. The origins of African Plasmodium vivax; insights from mitochondrial genome sequencing. PLoS One 2011; 6(12): e29137.

20 Miao M, Yang Z, Patch H, Huang Y, Escalante AA, Cui L. Plasmodium vivax populations revisited: mitochondrial genomes of temperate strains in Asia suggest ancient population expansion. BMC Evol Biol 2012; 12: 22.

21 Taylor JE, Pacheco MA, Bacon DJ, et al. The evolutionary history of Plasmodium vivax as inferred from mitochondrial genomes: parasite genetic diversity in the Americas. Mol Biol Evol 2013; 30(9): 2050–64.

22 Rodrigues PT, Alves JM, Santamaria AM, et al. Using mitochondrial genome sequences to track the origin of imported Plasmodium vivax infections diagnosed in the United States. Am J Trop Med Hyg 2014; 90(6): 1102–8.

23 Culleton R, Coban C, Zeyrek FY, et al. The origins of African Plasmodium vivax; insights from mitochondrial genome sequencing. PLoS One 2011; 6(12): e29137.

24 Wanderley DM, da Silva RA, de Andrade JC. Epidemiological aspects of malaria in the State of São Paulo, Brazil, 1983 to 1992. Rev Saude Publica 1994; 28(3): 192–7.

25 Curado I, Dos Santos Malafronte R, de Castro Duarte AM, Kirchgatter K, Branquinho MS, Bianchi Galati EA. Malaria epidemiology in low-endemicity areas of the Atlantic Forest in the Vale do Ribeira, São Paulo, Brazil. Acta Trop 2006; 100(1-2): 54–62.

26 Cerutti-Junior C, Boulos M, Coutinho AF, et al. Epidemiologic aspects of the malaria transmission cycle in an area of very low incidence in Brazil. Malaria Journal 2007; 6: 331–12.

27 Fonseca F. Plasmódio de primata do Brasil. Mem Inst Oswaldo Cruz 1951; 49: 543–551.

28 Garnham PCC. Malaria parasites and other haemosporidia, Blackwell Scientific Public, Oxford 1966. 1114 pp.

29 Singh B, Kim Sung L, Matusop A, et al. A large focus of naturally acquired Plasmodium knowlesi infections in human beings. Lancet 2004; 363(9414): 1017–24.

30 Deane LM, Deane MP, Ferreira Neto J. Studies on transmission of simian malaria and on the natural infection of man with Plasmodium simium in Brazil. Bull World Health Organ 1966; 35: 805–8.

